# Biomimetic Vasculatures by 3D-Printed Porous Molds

**DOI:** 10.1101/2021.09.27.461981

**Authors:** Terry Ching, Jyothsna Vasudevan, Shu-Yung Chang, Hsih Yin Tan, Chwee Teck Lim, Javier. G. Fernandez, Jun Jie Ng, Yi-Chin Toh, Michinao Hashimoto

## Abstract

Anatomically and biologically relevant vascular models are critical to progress our understanding of cardiovascular diseases (CVDs) that can lead to effective therapies. Despite advances in 3D bioprinting, recapitulating complex architectures (*i.e*., freestanding, branching, multilayered, perfusable) of a cell-laden vascular construct remains technically challenging, and the development of new techniques that can recapitulate both anatomical and biological features of blood vessels is of paramount importance. In this work, we introduce a unique, microfluidics-enabled molding technique that allows us to fabricate anatomically-relevant, cell-laden hydrogel vascular models. Our approach employed 3D-printed porous molds of poly(ethylene glycol) diacrylate (PEGDA) as templates to cast alginate-containing bioinks. Due to the porous and aqueous nature of the PEGDA mold, the calcium ion (Ca^2+^) was diffusively released to crosslink the bioinks to create hollow structures. Applying this technique, multiscale, multilayered vascular constructs that were freestanding and perfusable were readily fabricated using cell-compatible bioinks (*i.e*., alginate and gelatin methacryloyl (GelMA)). The bioinks were also readily customizable to either improve the compatibility with specific vascular cells or tune the mechanical modulus to mimic native blood vessels. Importantly, we successfully integrated smooth muscle cells and endothelial cells in a biomimetic organization within our vessel constructs and demonstrated a significant increase in monocyte adhesion upon stimulation with an inflammatory cytokine, tumor necrosis factor-alpha (TNF-α). We also demonstrated that the fabricated vessels were amenable for testing percutaneous coronary interventions (*i.e*., drug-eluting balloons and stents) under physiologically-relevant mechanical states, such as vessel stretching and bending. Overall, we introduce a versatile fabrication technique with multi-faceted possibilities of generating biomimetic vascular models that can benefit future research in mechanistic understanding of CVD progression and the development of therapeutic interventions.

## 1. Introduction

This paper describes the fabrication of freestanding, cell-laden vasculature models using 3D-printed porous molds to enable spatiotemporally controlled gelation of cell-laden hydrogels. Using our method of fabrication, we fabricated multiscale, multi-branching, multilayered vascular constructs that were freestanding and perfusable. Crucially, we demonstrated the incorporation of relevant vascular cells such as smooth muscle and endothelial cells in the multilayer arrangement. We highlighted the ability of the fabricated construct to test percutaneous coronary intervention (*i.e*., drug-eluting balloons and stents) under physiologically-relevant mechanical states, such as vessel stretching and bending. The versatility and diverse capabilities of our fabrication technique should allow us to biofabricate patient-specific vasculatures, which could aid future research in the mechanistic understanding of cardiovascular diseases (CVDs) and the development of therapeutic interventions.

CVDs remain to be one of the leading causes of mortality worldwide [1-3]. Despite the many advances in therapeutic interventions for CVDs, the pathophysiology of these diseases is still not fully understood due to the complex interplay between vessel anatomy, hemodynamic forces, and vascular cell responses [4]. Conventional experimental models in the form of 2D cell cultures and animal models are inadequate in providing sufficient insights into the development of CVDs in humans and their progression upon surgical or pharmacological interventions. 2D cellular models of a homogeneous population of cells cultured on a planar surface do not accurately recapitulate the anatomical arrangement of blood vessels, where stromal cells are found in the tunica media and the endothelium in the tunica intima layer [5, 6]. While animal models are indispensable in CVD research to recapitulate both the physical and biological attributes of a blood vessel comprehensively, they generally fall short as the underlying molecular, cellular, and physiological mechanism between animals and humans largely differs [7]. Furthermore, animal models are not suited for real-time visualization and monitoring of CVD progression (*e.g*., plaque accumulation and formation). Besides, the dimension and geometry of naturally occurring vasculatures are not precisely controlled, and some biophysical studies benefit from precisely engineered vasculature models. Therefore, there is a dire need for *in vitro* vascular models that can recapitulate human-specific anatomical vessel structures, flow dynamics, and the cellular environments to advance the understanding of CVD progression.

To date, researchers have employed various biofabrication strategies and technologies to improve the physiological relevance of vascular models in order to overcome the limitations of 2D cellular and animal models [8, 9]. Organ-on-chips (OoC) leveraged on microfluidic platforms to control the assembly of single or multiple cells in a microenvironment that aims to mimic *in vivo* organ-level function [7]. Due to their intrinsic perfusable nature, OoCs have been employed to model endothelialized vessels, hemodynamics, and vascular diseases (*e.g*., atherosclerosis, thrombosis, and stenosis) [10]. While vascular OoC systems continue to provide mechanistic insights into our understanding of vascular biology, there are still limitations to what OoCs can recapitulate [4, 11]. For instance, many OoCs still rely on rectangular channels embedded in polydimethylsiloxane (PDMS); such constructs fail to reconstruct the same geometry as blood vessels, which may alter the hemodynamics and the ensuing cellular responses. The confinement of the vasculature in the chip (as opposed to freestanding vasculatures) also limits the ability to replicate mechanical motions relevant to the vasculature.

To mimic the intricate geometries (*i.e*., bifurcations, curvatures) of blood vessels, additive manufacturing and, in particular, 3D bioprinting has been employed [12-15]. Namely, scaffold-free direct ink writing (DIW) [16-18], embedded bioprinting [19-22], light-assisted bioprinting [23], sacrificial molding [24-30], and coaxial bioprinting [31-34] have been employed to mimic the intricate architecture of vascular networks. Despite advances in bioprinting, the fabrication of complex vascular constructs remains a challenge because it involves the direct printing of bioinks, which must serve a dual function of supporting living cells and providing structural integrity to the vascular construct. Bioinks (*e.g*., extracellular matrix (ECM) and/or synthetic hydrogels) that can support living cells are usually soft after curing (*i.e*., the elastic modulus is <100 kPa), making them challenging to handle and print. Furthermore, a narrow range of thermal, mechanical, and chemical conditions must be met to ensure the printability of these bioinks and to prevent potential damage to integrated cells [21]. **Table S1-2** summarizes the currently available methods for fabricating vasculature structures in cell-laden hydrogels, highlighting their respective capabilities, advantages and limitations. These fabrication methods represent advances in 3D bioprinting, but the fabrication of complex vascular constructs to mimic their *in vivo* counterparts remains technically challenging to this day [35, 36].

The current constraints of 3D bioprinting motivated us to deviate from exclusively relying on direct 3D printing as an approach to fabricate vascular constructs. We took inspiration from the age-old molding technique, where the liquid material is shaped using a rigid mold with a hollow cavity that bears the shape of the desired design. However, unlike conventional molding processes where the filled liquid material solidifies *en masse*, we introduce a unique solidifying approach that permits controlled curing of bioinks to form tubular constructs. The porous molds were 3D-printed using high-resolution digital light processing (DLP) printers using poly(ethylene glycol) diacrylate (PEGDA) hydrogels. Controlled curing of bioinks was realized using 3D-printed porous molds permeated with crosslinker solution (*i.e*., a solution containing calcium chloride ion (Ca^2+^)). When the mold was filled with crosslinkable bioinks (*i.e*., sodium alginate), calcium ions diffused from the PEGDA molds into the bioink to prompt ionic crosslinking of the alginate-containing bioink from the region close to the wall of the mold. Importantly, the thickness of crosslinked alginate can be effectively controlled by varying the duration and concentration of the crosslinker solution. Uncrosslinked bioink was subsequently flushed out to realize a tubular construct that mimics blood vessels.

To that end, our proposed technique enables the fabrication of freestanding, multi-branching, and multilayered (*i.e*., up to four layers) vascular constructs in alginate-containing bioinks. The bioinks used for the formation of each layer in the hydrogel constructs can be customized by either adding bioactive materials to improve compatibility with relevant cells (*i.e*., gelatin methacryloyl (GelMA), fibrin, fibronectin) or adding synthetic materials to better mimic the mechanical properties of blood vessels (*i.e*., PEGDA). We leveraged on the laminar architectures of these hydrogel vascular constructs to culture smooth muscle cells and endothelial cells as concentric layers mimicking their physiological arrangements in a blood vessel. These cell-laden hydrogel constructs could be maintained for up to 10 days under perfusion culture. To highlight that the endothelium in the vascular construct was functional, we activated the endothelial cells with pro-inflammatory cytokines and observed an increase in monocyte adhesion. Lastly, we demonstrated the capability of these vascular constructs to mimic the cyclical movement of the coronary artery and the insertion of percutaneous coronary interventions (*i.e*., drug-eluting balloon, drug-eluting stent), showcasing the possibility for future development of vascular models for coronary disease research. Our method enabled the fabrication of biomimetic vascular models with anatomically and biological accuracy, which shall serve as models to further our understanding of CVD pathophysiology and testbeds for future therapeutic interventions.

## 2. Materials and Methods

### 2.1. Materials

Sodium alginate (alginic acid sodium salt from brown algae, low viscosity), calcium chloride (CaCl_2_), fibrinogen (from bovine plasma), thrombin (from bovine plasma), poly(ethylene glycol) diacrylate (PEGDA) (molecular weight, MW = 700), quinoline yellow (QY) were purchased from Sigma-Aldridge (St. Louis, MO, USA). Fibronectin (human), ruthenium (Ru), sodium persulfate (SPS) was purchased from Advanced Biomatrix (San Diego, CA, USA). Lyophilized GelMA and lithium phenyl-2,4,6-trimethylbenzoylphosphinate (LAP) were purchased from Cellink (Boston, MA, USA). Rabbit VE-Cadherin, and mouse CD31 primary antibodies were purchased from Cell Signaling Technology (Danvers, MA, USA). Rabbit and mouse anti-α-smooth muscle actin (α-SMA) primary antibodies were purchased from Abcam (Cambridge, MA, USA). HEPES buffer solution, SYTOX™ green nucleic acid stain, fetal bovine serum (FBS), paraformaldehyde (PFA) (16 % (w/v)), and all secondary antibodies were purchased from Thermo Fisher Scientific (Waltham, MA, USA). Recombinant Human TNF-alpha was purchased from R & D Systems (Minneapolis, MN, USA). RapiClear^®^ was purchased from SUNJin Lab (Hsinchu City, Taiwan). Rubber washer O-rings (for watch crown), PTFE tubing were purchased from Aliexpress (Hangzhou, China). Drug-eluting balloons (Mini Trek, diameter = 2 mm, length = 20 mm) and drug-eluting stents (Xience Alpine, diameter = 2 mm diameter, length = 8 mm) were provided by Abbott (Chicago, IL, USA).

### 2.2. Fabrication and preparation of PEGDA molds

14 % (v/v) PEGDA (MW = 700) was prepared in 10 mM HEPES buffer solution. Ru and SPS were added as photoinitator at a final concentration of 0.2 and 2 mM, respectively. QY was added as photoabsorber at a final concentration of 0.5 mg/mL. Computer-aided design (CAD) of the molds was designed in Rhinoceros (Robert McNeel & Associates, WA, USA). Asiga Pico 2HD (Sydney, Australia) was employed for all 3D-printed parts. The printed PEGDA molds were soaked in deionized (DI) water overnight to leech out the remaining photoabsorber. For sterilization, the PEGDA parts were sterilized by soaking in 70 % ethanol for 1 hr. Subsequently, the PEGDA parts were soaked in three changes of HEPES buffer solution, 1 hr each, to remove the residual ethanol. The PEGDA parts were kept in 0.2 M CaCl_2_ solution at 4°C until use. Before usage, the PEGDA parts were fitted into a two-part, 3D-printed rigid shell, as demonstrated in earlier works [37]. The two-part rigid shell functioned as a support frame around the soft PEGDA molds (**Fig. 1A**). Binder clips were used to maintain conformal contact between the rigid shells to ensure conformal contact (**Fig. 1C, Movie S4**). The rigid shells were fabricated using a combination of laser-cut poly(methyl methacrylate) (PMMA) sheets and 3D-printed components (Dental SG resin, Formlabs, MA, USA).

**Figure 1.**
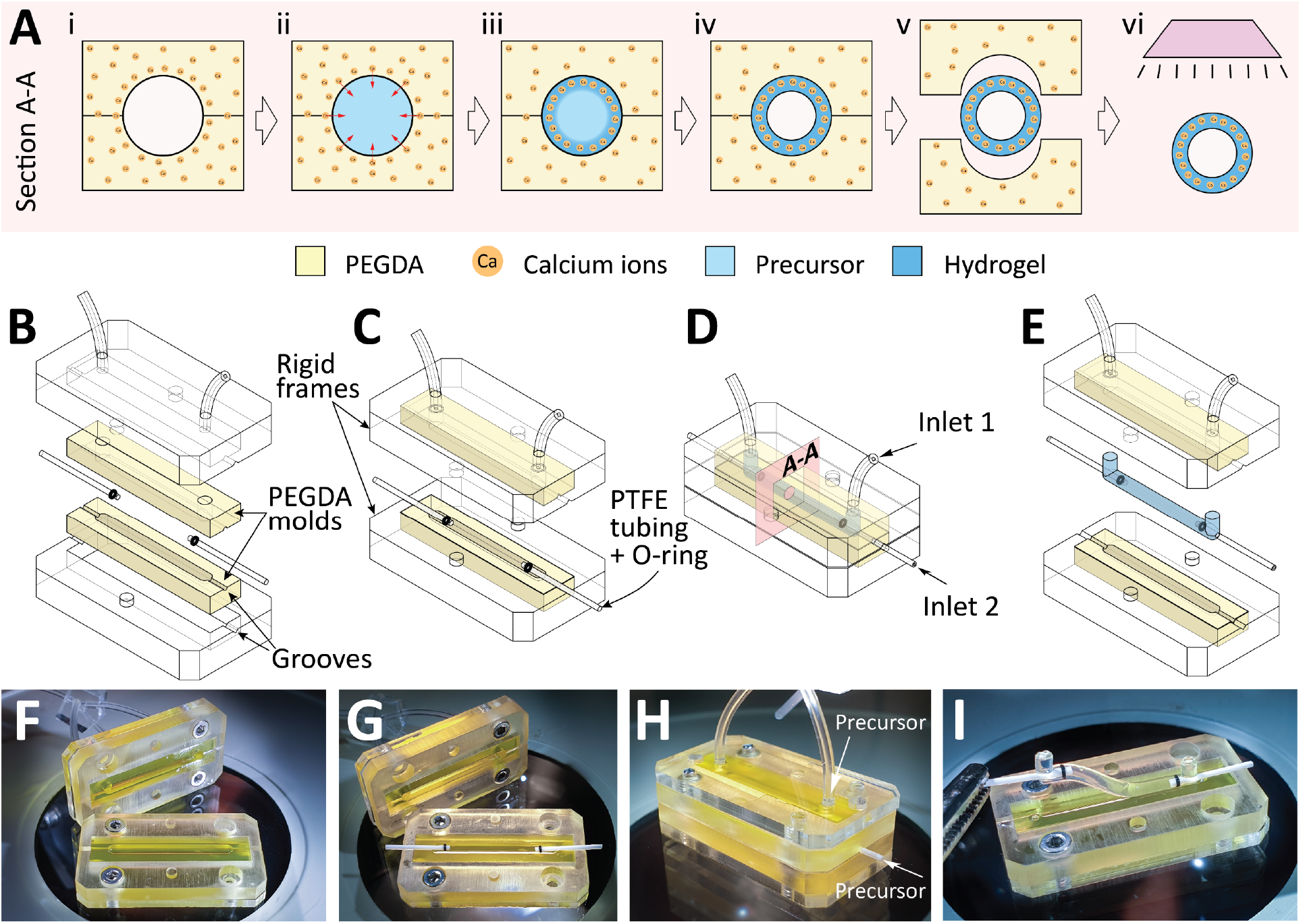
Fabrication of free-standing hydrogel vasculatures using 3D printed porous mold. **A)** Cross-sectional (A-A) illustrations of the steps for the fabrication of hollow tubular hydrogels. Hydrogel-based bioinks are crosslinked in the 3D printed PEGDA mold, followed by photopolymerization. **B-E)** 3D schematics illustrating the step-by-step fabrication of freestanding vascular constructs. **F-I)** Photographs showing the step-by-step fabrication of vascular constructs.

### 2.3. Preparation of different precursor bioinks

The precursor bioink containing alginate and GelMA (Alg-GM) to fabricate cell-free hydrogel vascular constructs were prepared by weighing the desired amounts of lyophilized GelMA, LAP, and alginate and reconstituting them in 10 mM HEPES buffer solution. The precursor bioinks employed for cell-laden vascular constructs were prepared by weighing the desired amounts of lyophilized GelMA, LAP, and alginate and reconstituting them in a solution containing 80 % (v/v) HEPES (10 mM), 10 % (v/v) FBS, and 10 % (v/v) fibronectin solution (0.5 mg/mL stock solution). The precursor underwent constant stirring at 45°C for 1 hr. The mixtures were prepared on the day of the experiment and were kept at 37°C until use. Cells were mixed with the precursor prior to the fabrication of freestanding vascular constructs.

To fabricate a two-layer vascular construct consisting of a fibrin ECM layer (**Fig. 4F, Fig. S2**), the first precursor contained 2 % (w/v) alginate, 6.7 % (w/v) GelMA, 0.25 % (w/v) LAP, and 50 U/mL thrombin prepared in HEPES buffer (10 mM) that underwent constant stirring at 45°C for 1 hr. The second precursor solution contained 15 mg/mL fibrinogen, 6.7 % (w/v) GelMA, and 0.25 % (w/v) LAP prepared in HEPES buffer (10 mM) that underwent constant stirring at 45°C for 1 hr.

### 2.4. Mechanical properties of different bioink formulations

To characterize the bulk mechanical properties of various bioinks, pre-polymer solutions with varying compositions (e.g., alginate (2 % (w/v)), GelMA (7 % (w/v)), PEGDA (5 – 20 % (w/v)) and LAP (0.25 % (w/v))) were pipetted to a PDMS mold with a cylindrical cavity (8 mm in diameter, 2.6 mm in height). Subsequently, the molds were exposed to ultra-violet (UV) at 405 nm for 15 s. Next, the cylindrical samples were soaked in 0.2 M CaCl_2_ solution for 15 min, followed by HEPES buffer solution overnight before the compression test. Compression tests were carried out using the TA ElectroForce 3230-ES dynamic mechanical test instrument (TA Instruments, New Castle, DE, USA). Tests were carried out with a ramp force rate of 10.0 N·min^-1^ up to 18 N at room temperature. The compressive modulus was determined as the slope of the linear region corresponding with 0 – 10 % strain [31, 38].

### 2.5. Fabrication of freestanding hydrogel vascular constructs

3D printed PEGDA molds pre-soaked in CaCl_2_ solution were first assembled with the supporting rigid frames, PTFE tubes (outer diameter = 0.77 mm), and o-rings (1.4 mm × 0.6 mm × 0.4 mm) (**Fig. 1A**). All the parts were secured in place using a binder clip (**Movie S4**). Inlets 1 and 2 were coupled together using a PDMS Y-channel and connected to a syringe containing the Alg-GM bioink precursor. The Alg-GM precursor was perfused into the mold at 100 µL·min^-1^ to fill the mold cavity completely. Upon filling the cavity, the flowrate was changed to 10 µL·min^-1^ and was held at that flowrate for the desired duration (*i.e*., dwell time), during which Ca^2+^ ions in the porous PEGDA molds would diffuse into the cavity and crosslink the alginate fraction in the precursor bioink. After which, HEPES buffer was perfused through the mold to remove the uncrosslinked Alg-GM precursor. The freestanding hydrogel network was then removed from the mold and was exposed to UV light (Anycubic Cure Machine, Shenzhen, China) for 15 s to crosslink the embedded GelMA.

### 2.6. Microfluidic setup to characterize alginate crosslinking dynamics

A parallel plate microfluidic device was constructed to characterize the crosslinking dynamics of Alg-GM bioinks by Ca^2+^ ions released by porous PEGDA blocks (**Fig. 2**). The microfluidic device consisted of a PDMS sheet (thickness = 0.8 mm) sandwiched between two laser-cut PMMA sheets. Through holes were included on the PMMA sheets that functioned as inlet ports, outlet ports, and holes for mechanical fasteners (M2 screws). During assembly, the PEGDA block, which was pre-soaked in CaCl_2_ solution, was placed in the microfluidic device and sealed using the PMMA sheets. The device was mounted onto a fluorescence microscope, and Alg-GM precursor solution spiked with 10 µm fluorescence particles (Fluor Polystyrene, Magsphere, Pasadena, CA, USA) was perfused through the inlet port using a syringe pump. Movement of particles was recorded and analyzed by particle image velocimetry (PIV) using a toolbox developed by Thielicke et al. on Matlab (Natick, MA, USA) named PIVlab [39-41].

**Figure 2.**
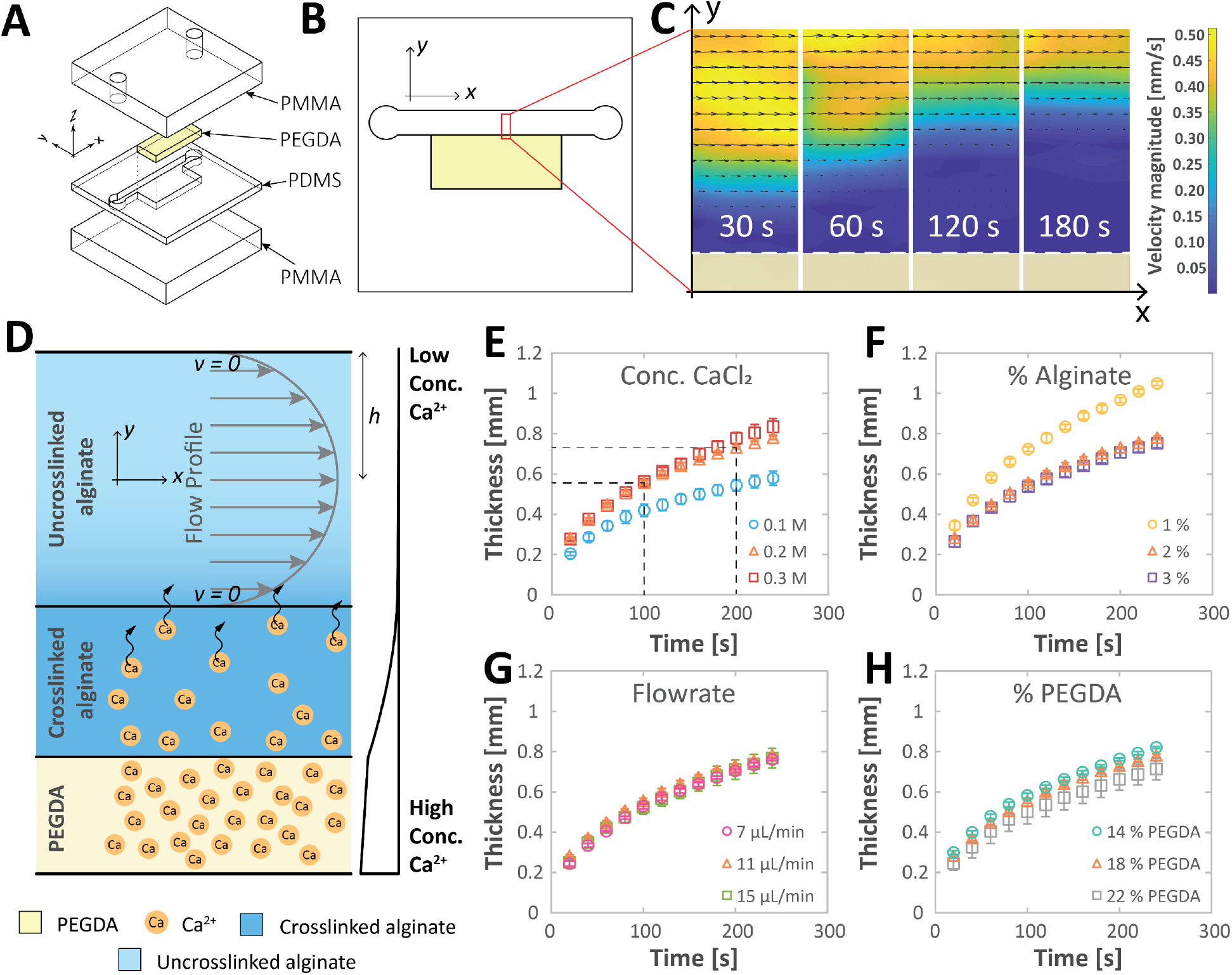
Dynamic crosslinking of alginate-based bioink by the diffusion of calcium ions diffused from a PEGDA block. Schematic illustrations of the microfluidic device used to characterize the crosslinking process: **A)** exploded view and **B)** top view. **C)** Particle image velocimetry (PIV) analysis of the precursor solution upon the perfusion through the microfluidic device at different time points (30 s, 60 s, 120 s, and 180 s). White dashed line denotes the boundary between the PEGDA block and precursor bioink. **D)** Graphical schematic illustrating the diffusion of calcium ions across a concentration gradient inside the microfluidic device. Plots showing the thickness of crosslinked alginate over the dwell time when varying the following parameters: **E)** molar concentration of CaCl2, **F)** % (w/v) of alginate in precursor solution, **G)** flowrate of the precursor solution (bioinks), and **G)** % of PEGDA (v/v) employed to fabricate the PEGDA block.

### 2.7. Fabrication and maintenance of cell-laden vascular constructs

#### 2.7.1. Cell culture and maintenance

Primary Human Umbilical Vein Endothelial Cells (HUVEC) (Lonza, Basel, Switzerland) were routinely cultured in a 37°C, 5 % CO_2_ incubator. Cells were supplemented with EGM−-2 endothelial cell growth medium (Lonza, Basel, Switzerland). Cells were passage at 70 – 80 % confluency, and only cells between passage 3-10 were used for experiments. Primary human aortic smooth muscle cells (SMCs) (Lonza, Basel, Switzerland) were routinely cultured in a 37°C, 5 % CO_2_ incubator in SmGM− growth medium (Lonza, Basel, Switzerland). The SMCs were passaged at 70 – 80 % confluency, and only cells with the passage 3 – 10 were used for experiments.

#### 2.7.2. Fabrication and setup of cell-laden vascular constructs

To create SMC-laden vascular constructs, SMCs were dissociated from the culture flask and centrifuged to form a cell pellet. The supernatant media was removed, and precursor solution (2 % (w/v) alginate, 6.7 % (w/v) GM, 0.25 % (w/v) LAP, 10 % (v/v) FBS, 50 μg/mL fibronectin) was added to the cell pellet and resuspended by gentle pipetting to form a homogenous cell suspension (8 × 10^6^ cells mL^-1^). The SMC suspension was loaded into a 3 mL syringe and fitted with a 22G, blunt dispenser needle. After fabricating the vascular construct using the precursor bioink containing the SMCs as described in section 2.5 (**Movie S4**), the freestanding vascular construct was connected to a 3D printed frame and coupled to peristaltic pumps as described in our earlier work [37]. After seven days of culture under continuous perfusion culture at a flowrate of 2.2 µL·min^-1^, the vascular construct was seeded with HUVECs. 30 μL of HUVECs suspended in EGM−-2 endothelial cell growth medium at a density of 5 × 10^6^ cells mL^-1^ were perfused by hand into the vascular construct. The vascular construct was incubated for 30 min to facilitate cell adhesion. After 30 min, the vascular construct was flipped 180°, perfused with a new batch of HUVEC cell suspension, and incubated for another 30 mins. Next, the vascular construct was reconnected to the peristaltic pump to commence continuous perfusion culture at ∼1.1 µL·min^-1^ (**Fig S3**). Peristaltic pumps were fabricated as previously described [42].

#### 2.7.3. Monocyte adhesion assay

Monocyte adhesion assay was performed three days after seeding HUVECs. Vascular constructs were treated with TNF-α (10 ng/mL) for 4 hr under constant perfusion with a peristaltic pump (∼1.1 µL·min^- 1^). After 4 hr, the vascular constructs were perfused with regular EGM−-2 medium for 15 min. Vascular constructs were then perfused with U937 monocytes (pre-labeled with CellTracker− Green at a density of 1 × 10^6^ cells mL^-1^ for 30 min. Next, the vascular constructs underwent three washes with 1*×* Dulbecco’s phosphate-buffered saline (D-PBS) (Nacalai Tesque, Kyoto Japan) and were fixed for whole-mount imaging under a fluorescence microscope (Zeiss Observer D1, Oberkochen, Germany). The number of monocytes adhering to the vessel wall was quantified by cell counting analysis performed using ImageJ [43].

#### 2.7.4. Imaging and analysis

Immunostaining and confocal microscopy (Zeiss LSM 710, Oberkochen, Germany) were used to assess the vascular constructs. The vascular constructs were first washed with 1*×* PBS for several minutes. Next, 4% PFA was perfused through the vascular constructs while submerged in the same 4% PFA solution for 40 min. The vascular constructs were washed with 1*×* PBS (perfused and submerged) three times, 15 minutes each. Next, the vascular constructs were permeabilized (1% Triton-X-100) for one day at 4°C and blocked (2% BSA and 1% Triton-X-100) for one day at 4°C. The vascular constructs were incubated with primary antibodies of interest at a concentration of 1:200 for two days in a solution of 2% BSA and 0.2% Triton-X-100 at 4°C (**Table S3**). The unbounded primary antibodies were removed by first washing the vascular constructs with washing buffer (0.1% Triton-X-100 in 1*×* PBS) twice at room temperature, 1 hr each. The vascular constructs were then kept in washing buffer on an orbital shaker at 4°C overnight. Next, secondary antibodies were incubated at a concentration of 1:500 with the vascular construct for two days in a solution of 2% BSA and 0.2% Triton-X-100 at 4°C. SYTOX−-green was also added at a concentration of 1:10,000 in this secondary antibody step. The unbounded secondary antibodies were removed by first washing the vascular constructs with washing buffer twice at room temperature, 1 hr each, and kept in washing buffer on an orbital shaker at 4°C overnight. After washing, the vascular constructs were rinsed with PBS three times, 30 min each. Lastly, the vascular constructs were kept in RapiClear^®^ gel for one day prior to confocal imaging.

### 2.8. Setup and evaluation of mechanically stretchable branching vascular constructs

The expandable balloon to mimic the beating heart movement was fabricated by first casting a thin film (∼0.5 mm thickness) of Ecoflex™ 00-50 (Smooth-On, Macungie, PA, USA) in a petri dish (100 mm diameter). The periphery of the Ecoflex thin film was then permanently sealed to a PMMA sheet using silicon sealant. A hole was drilled to the middle of the PMMA sheet for coupling of tubing. A syringe pump was employed to inflate and deflate the Ecoflex balloon. The extent of inflation and deflation was controlled by varying the displacement of the syringe plunger. The expandable balloon with branching vascular construct was mounted to a fluorescence microscope (Zeiss Observer D1, Oberkochen, Germany) to visualize the flow of tracer particles. Yellow-green fluorescence tracer particles of 10 µm (Fluor Polystyrene, Magsphere, Pasadena, CA, USA) were diluted to a concentration of 1 × 10^7^ particles mL^-1^ and perfused through the vascular construct with a syringe pump at a rate of 2 µL·min^-1^. Movement of particles was recorded, and particle image velocimetry (PIV) analysis was performed using a toolbox developed by Thielicke et al. on Matlab (Natick, MA, USA) named PIVlab [39-41].

### 2.9. Insertion of drug-eluting balloon and stent

For the vascular constructs employed for the insertion of the drug-eluting balloon, the PEGDA mold was designed with a diameter of 2.4 mm, and the constriction was designed with a diameter of 1.46 mm. For the vascular construct employed for the insertion of the drug-eluting stent, the diameter of the PEGDA mold was designed with a diameter of 2.8 mm and the constriction diameter of 2.17 mm. Precursor bioink containing 2% (w/v) alginate, 6.7% (w/v) GelMA and, 0.25% (w/v) LAP were used to fabricate the vascular constructs. Dwell time of 1 min was used, resulting in a wall thickness of ∼550 µm. The fabricated vascular constructs were mounted on a 3D printed frame. Insertion of balloon and stent was performed following the instructions by the manufacturer.

## 3. Results and Discussion

### 3.1. Experimental Design

We aimed to develop a method to fabricate biomimetic vascular constructs by simultaneously taking advantage of two fabrication methods: molding and coaxial bioprinting. In favor of fabricating freestanding vascular constructs to mimic blood vessels with complexity, we explored the use of a negative mold approach (**Fig. S1**). However, the negative mold approach was not suitable for generating hollow constructs, and we then took inspiration from existing coaxial bioprinting techniques: dynamic crosslinking between two sheath flows of a crosslinkable bioink (*e.g*., sodium alginate) and the crosslinker solution (*e.g*., CaCl_2_) within the mold. Unlike coaxial bioprinting, we did not employ multiple coaxial nozzles to dispense concentric streams of crosslinker and crosslinkable bioinks. Instead, we relied on 3D-printed porous molds that were permeated with the crosslinker solution (**Fig. 1A-i**). These molds were 3D-printed as a two-part entity, with a negative cavity in the shape of the desired vascular network. As previously demonstrated using a digital light processing printer, we printed biomimetic vascular networks in PEGDA hydrogels [37]. Here, we specifically utilized high molecular weight PEGDA (MW = 700) to ensure that the PEGDA mold was sufficiently porous so that Ca^2+^ ions can be efficiently contained in the mold when soaked in the solution of CaC1_2_. Permeability of PEGDA hydrogels (MW = 700) were studied and reported in existing literature [44-46]. The advantage of using a porous mold is the capability of fabricating hollow constructs. As crosslinkable precursor bioink (*i.e*., alginate-containing bioink) was perfused into the porous mold, Ca^2+^ ions diffused radially into the mold cavity and prompted ionic crosslinking of alginate (as demarcated by red arrows in **Fig. 1A-ii, iii**). The crosslinking reaction was discontinued to create hollow vascular constructs by perfusing HEPES buffer solution to evacuate the uncrosslinked precursor bioink (**Fig. 1A-iv**). The hollow vascular construct was then removed from the mold and exposed to ultra-violet (UV) light to photocrosslink another biomaterial (*i.e*., GelMA) present in the hydrogel (**Fig. 1A-v, vi**).

**Fig. 1B-I** illustrate the step-by-step fabrication of freestanding vascular constructs. Rigid frames were fabricated using a combination of 3D printing and laser-cut PMMA sheets. The rigid frames aided to ensure conformal contact between the two-part PEGDA mold (**Fig. 1B-C, F-G**). PTFE tubings were incorporated directly into the fabrication process, by including grooves on the mold, to eliminate the hassle of connecting tubings for subsequent perfusion culture (**Fig. 1B**). O-rings were also included on the PTFE tubings to provide an effective seal between the hydrogel and PTFE tubings (**Fig. 1C**). After perfusion of precursor solution and flushing with HEPES buffer solution (**Fig. 1A, D, H**), the PTFE tubings were directly integrated with the vascular constructs for easy handling (*i.e*., with a tweezer) and performing perfusion culture (**Fig. 1E, I**).

### 3.2. Control and characterization of alginate crosslinking mechanism by PEGDA molds

To determine the crosslinking characteristics of alginate-containing bioinks when perfused into the PEGDA mold, we devised a simplified microfluidic setup. A parallel plate flow chamber was fabricated to visualize the crosslinking of sodium alginate solution when flowing over a porous PEGDA matrix permeated with calcium ions (**Fig. 2A-B**). Fluorescent tracer particles were mixed in with the alginate precursor solution to visualize the flow profiles of the precursor solution and the subsequent crosslinking into the gel phase. When the precursor was perfused into the channel at a constant flowrate (*x*-direction), we observed that the fluorescent beads close to the PEGDA interface were instantly immobilized upon contact, indicating that they were trapped in the crosslinked alginate matrix (**Movie S5**). As time progresses, the layer of immobilized beads grew thicker in the *y*-direction (**Fig. 2C**, dark-blue area).

The thickness of crosslinked alginate is dependent on the distance traveled by the Ca^2+^ ions that were primarily driven by diffusion across a concentration gradient. Fick’s first law of diffusion describes the diffusion of molecules across a concentration gradient:

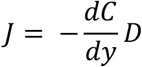

Where *J* is the diffusion flux (*i.e*., the net rate of particle/molecules moving through an area), 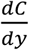 is the concentration gradient and, *D* is the diffusion coefficient or diffusivity (*i.e*., the diffusion coefficient of Ca^2+^ ions in alginate hydrogel). When uncrosslinked alginate precursor contacted the PEGDA hydrogel that had a high concentration of Ca^2+^ ions, the concentration gradient resulted in a net movement of Ca^2+^ ions in the *y*-direction (**Fig. 2D**). The Ca^2+^ ions prompted rapid ionic crosslinking of alginate monomer to form calcium alginate hydrogel. The concentration gradient continues to drive the diffusion of Ca^2+^ ions over time, increasing the thickness of crosslinked alginate hydrogels. The mean distance traveled of any given molecule in a solution is usually estimated using the Einstein-Smoluchwski relation [47]:

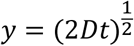

Where *y* is diffusing solute in one direction (y-direction) after elapsed time *t*, (*i.e*., diffused distance of the calcium ions), *D* is the diffusion coefficient of a solute in a given medium, and *t* is the elapsed time since diffusion began. It should be noted that the Einstein-Smoluchwski relation can only provide an approximation. Nevertheless, it predicts the general trend we observed in our experiments. The experimental plots representing the diffusion of Ca^2+^ ions across the channel length (*y*) followed the trend described by the Einstein-Smoluchowski relation, where the thickness of the crosslinked alginate was proportional to t^1/2^ (**Fig. 2E-H**).

Next, we varied four separate parameters of the inks and flows to study the rate of alginate crosslinking in the *y*-direction. First, we varied the concentration of calcium chloride that the PEGDA molds (*i.e*., 0.1, 0.2, 0.3 M) were pre-soaked in. Secondly, we varied the concentration of the alginate in the perfused precursor solution (*i.e*., 1, 2, 3 % (w/v)). Thirdly, we varied the flowrates at which the precursor solutions were perfused through the microfluidic device (*i.e*., 7, 11, 15 μL/min). Lastly, we varied the PEGDA concentration employed during 3D printing the PEGDA block (*i.e*., 14, 18, 22 % (v/v)) for the variation of the porosity of the molds. The results of the study were summarized (**Fig. 2E-H**). The plot described by the orange marker in all four graphs represented the thickness of immobilized beads over time t with the following parameters: (1) 0.2 M CaCl_2_, (2) 2 % alginate, (3) 11 μL/min, and (4) 18 % PEGDA.

Experimentally, we found that the key parameters to vary the rate of diffusion of calcium ions were (1) concentration of CaCl_2_ and (2) % of alginate. The higher concentration of CaCl_2_ led to the faster diffusion rate of calcium ions because the higher concentration gradient of calcium ions resulted in the larger diffusion flux *JJ* (**Fig. 2E**). We observed that the lower % alginate led to the more rapid rate of diffusion distance, presumably because the resulting hydrogel with the lower % alginate was more porous with the larger effective diffusion coefficient *D* (**Fig. 2F**). Next, we explored the effect of the flowrate on the formation of the alginate hydrogels. Interestingly, we found that varying the flowrate did not result in a noticeable change in the formation of the alginate hydrogels (**Fig. 2G**). This phenomenon can be attributed to the no-slip boundary condition where the flow distribution *v*(*y*) of viscous flow between parallel plates is described as:

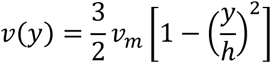

And mean velocity *v*_*m*_ is:

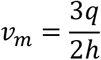

where *q* is the volumetric flowrate, and *h* is the distance from the channel wall to the midway point (y = 0) (**Fig. 2D**). At the fluid-solid boundaries (y = *h*), *v*(y) is equal to zero (see also **Movie S5**). As such, at any given time, the infinitesimal layer of uncured precursor on the fluid-solid boundaries was assumed to have zero velocity; thus diffusion of Ca^2+^ ions was not affected by the different flowrate. Lastly, we postulated that the higher % of the PEGDA block would result in a matrix with the lower porosity and the higher diffusion rate of Ca^2+^ ions. However, experimentally, we observed little difference in the formation of the alginate hydrogels with the variation of % PEGDA in the block (**Fig. 2H**). High % PEGDA (*i.e*., 18 % and 22 %) were also brittle and unsuitable for subsequent experiments. These experiments demonstrated that varying the concentration of CaCl2 solution and % of alginate could substantially alter the rate of crosslinking of alginate hydrogels. Varying the flowrate of precursor solution and % PEGDA of printed blocks did not alter the wall thickness substantially, however. Crucially, we found that altering the residence time (or dwell time) of the precursor inside the porous mold prior to the evacuation of uncrosslinked precursor was an effective and consistent parameter to precisely control the thickness of cured alginate, as shown by the small error bars (defined as standard deviation, N = 3). For example, using the parameters of the orange scatter plot, a dwell time of 100 s resulted in a thickness of 0.56 mm, and a dwell time of 200 s resulted in a thickness of 0.73 mm (see dashed lines in **Fig. 2E**). Experimentally, the slow growth of the alginate hydrogels gives a wide operation window to control the thickness of the formed alginate hydrogels. These tests provided information that guided the subsequent experiments.

### 3.3. Customization of bioink properties

Next, we evaluated the feasibility of fabricating vascular constructs using bioinks of different formulations to tune their mechanical and cell support properties. While the alginate hydrogel is known to be noncytotoxic, it does not support cell spreading, motility, and proliferation [48]. Therefore, we explored blending other bioactive materials with alginate to form the bioinks to improve compatibility with relevant vascular cells. GelMA is one such candidate that has been commonly blended with alginate to fabricate tubular constructs using coaxial bioprinting techniques [31, 32]. GelMA has been shown to improve cell spreading, motility, and proliferation [49, 50]. We tested different formulations of bioinks that contained 1 – 3 % (w/v) alginate and 2.5 – 15 % (w/v) GelMA (**Fig. 3A**). We found that a low concentration of alginate content (*i.e*., 1 % (w/v)) was not feasible in fabricating the vascular constructs; the low alginate concentration resulted in a mechanically weak hydrogel that collapsed under its weight, making them difficult to handle when removing from the PEGDA molds. A high concentration of GelMA (≥ 10 % (w/v)) also proved to be challenging to handle as GelMA was susceptible to thermal gelation at room temperature (**Fig. 3A**, yellow portion). When both the concentrations of alginate and GelMA were high (**Fig. 3A**, green portion), the viscosity of the bioink precursor was high and was prone to cause leakages when perfused through the PEGDA mold. Overall, we found the condition as highlighted in red favorable when fabricating vascular constructs using the PEGDA molds.

**Figure 3.**
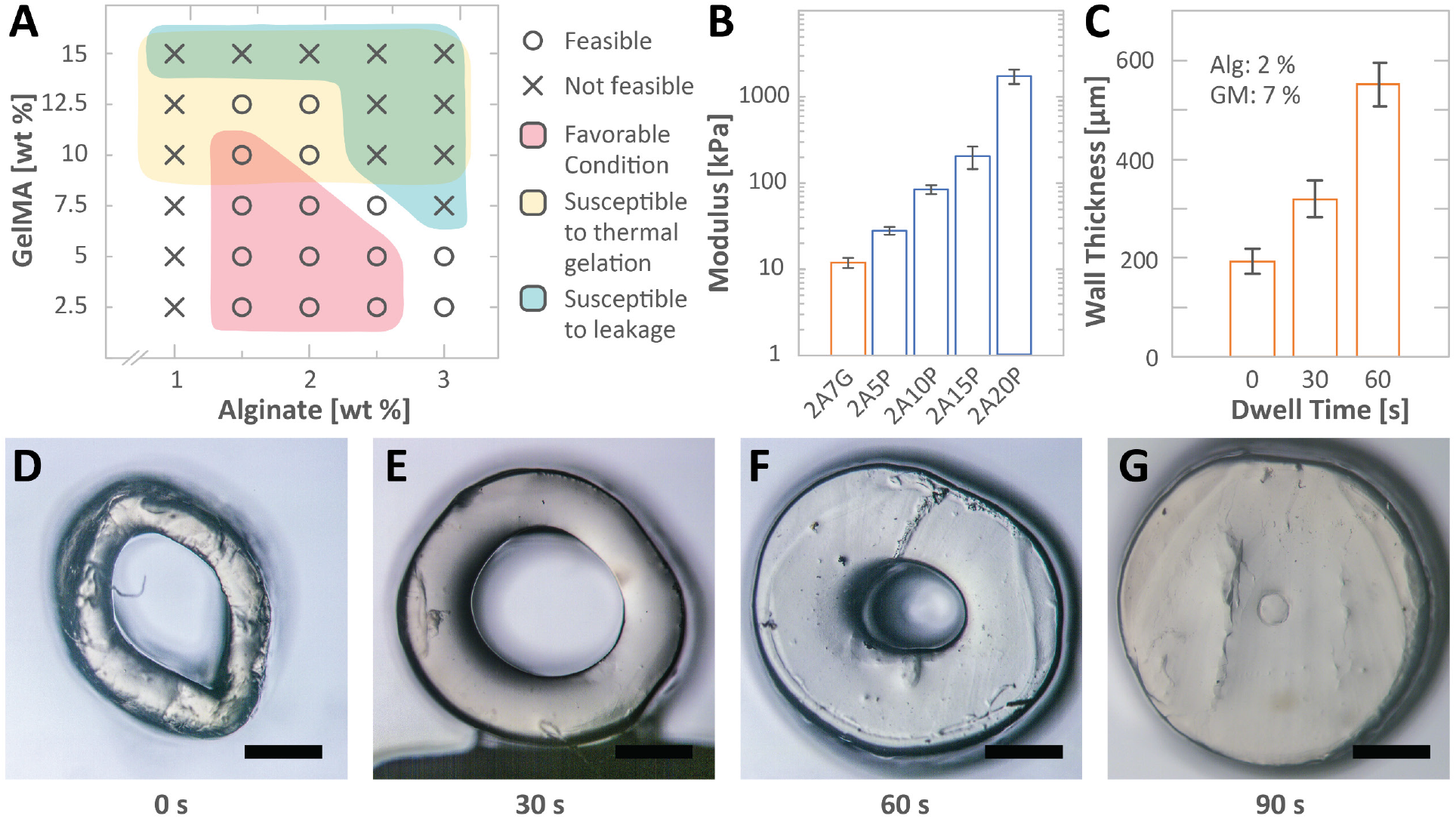
Characerization of bioinks and control over the wall thickness of fabricated vascular constructs. **A)** Phase diagram showing the feasibility of bioinks with varying proportions of alginate and GelMA to fabricate the hydrogel vascular constructs. **B)** Bar graph showing the elastic modulus of hydrogel with different formulations (*e.g*., 2A: 2 % (w/v) alginate, 7G: 7 % (w/v) GelMA, 5P: 5 % (v/v) PEGDA; the same nomenculture applies to describe the materials consisting of alginate, GelMa and PEGDA). **C)** Bar graph showing the relationship between the wall thickness and the dwell time for 2A7G. **D-G)** Micrographs showing the cross-sections of the vascular constructs with the different dwell times. Scale bars: 500 µm.

We also explored the use of other bioinks that tuned the overall stiffness of the vascular constructs. While GelMA contains cell-adhering moieties that supported the growth of relevant cell types, it does not have the mechanical properties that match *in vivo* blood vessels. For instance, various arteries in the human body were reported to have elastic moduli of 0.1 – 1.5 MPa [51-54]. However, the measured elastic modulus of Alg-GM (2A7G: 2 % (w/v) Alg, 7 % (w/v) GM) was ∼12 kPa, which was not in the order of the blood vessels *in vivo* (**Fig. 3B**). The addition of stiff materials (such as PEGDA) allows tuning the mechanical stiffness of hydrogels. For example, a blend consisting of 2 % (w/v) of alginate and 5% (v/v) of PEGDA (denoted as 2A5P) had an elastic modulus of 28 kPa. Increasing the PEGDA concentration from 5% to 20% (v/v) could increase the elastic modulus by up to 60-fold (∼ 1.7 MPa) (**Fig. 3B**). This experiment verified the suitability of using PEGDA as a candidate to enhance the mechanical stiffness of alginate-containing bioinks. The blending of PEGDA with alginate allowed the fabrication of vascular construct with elastic modulus comparable to *in vivo* blood vessels (> 100 kPa). The added stiffness also facilitates the handling of the vascular constructs.

Lastly, we demonstrated that our method effectively controlled the thickness of the fabricated vascular constructs using relevant bioinks (*i.e*., Alg-GM). We varied the dwell time and measured the wall thickness of the fabricated hydrogel vascular constructs (**Fig. 3B, D-G**). We employed a bioink blend of 2 % (w/v) alginate and 7 % (w/v) GelMA. By varying the dwell time up to 1 min, the wall thickness was tuned to be ∼200 – 550 µm (**Fig. 3C**). Further increase in the dwell time (1 min 30 s) resulted in the clogging of the channel (**Fig. 3G**). Effective thickness control should enable end-users to tailor the scale and thickness of vascular constructs for increased physiological relevance.

### 3.4. Fabrication of anatomically-relevant vascular constructs

Applying our fabrication technique, we demonstrated the fabrication of anatomically-relevant vascular constructs (*i.e*., freestanding, branching, multilayered, and perfusable). The design of the 3D-printed PEGDA mold determines the architecture of the fabricated vasculature. We fabricated a single-channel vascular construct (**Fig. 4A**), a bifurcating vascular construct (**Fig. 4B**), and a hierarchical, multi-branching vascular construct (**Fig. 4C**). In addition to varying architecture, blood vessels are composed of endothelial cells (ECs), smooth muscle cells (SMCs), fibroblasts, and extracellular matrix (ECM) in a multilayered configuration [55]. Advantageously, our fabrication technique allows the biomimicry of such multilayered configurations. By successively perfusing different batches of bioinks into the mold, we fabricated a bifurcating vascular construct with four laminar layers (**Fig. 4D**), as highlighted by the different colored fluorescence beads encapsulated within each layer (**Fig. 4E**). One key benefit of the multilayered architecture is the ability to customize and tune the biochemical composition of each Alg-GM hydrogel layer. For instance, the ability to customize the biochemical composition is beneficial when incorporating different vascular cells into the constructs. To demonstrate the ability to include other cell-responsive ECMs, we used fibrinogen and thrombin to create a layer of fibrin ECM (**Fig. 4F, S2, Movie S6**). While the demonstration in this work was limited to GelMA and fibrin, we anticipate the possibility of employing other ECM blends such as collagen methacrylate (ColMA) and hyaluronic acid methacrylate (HAMA).

**Figure 4.**
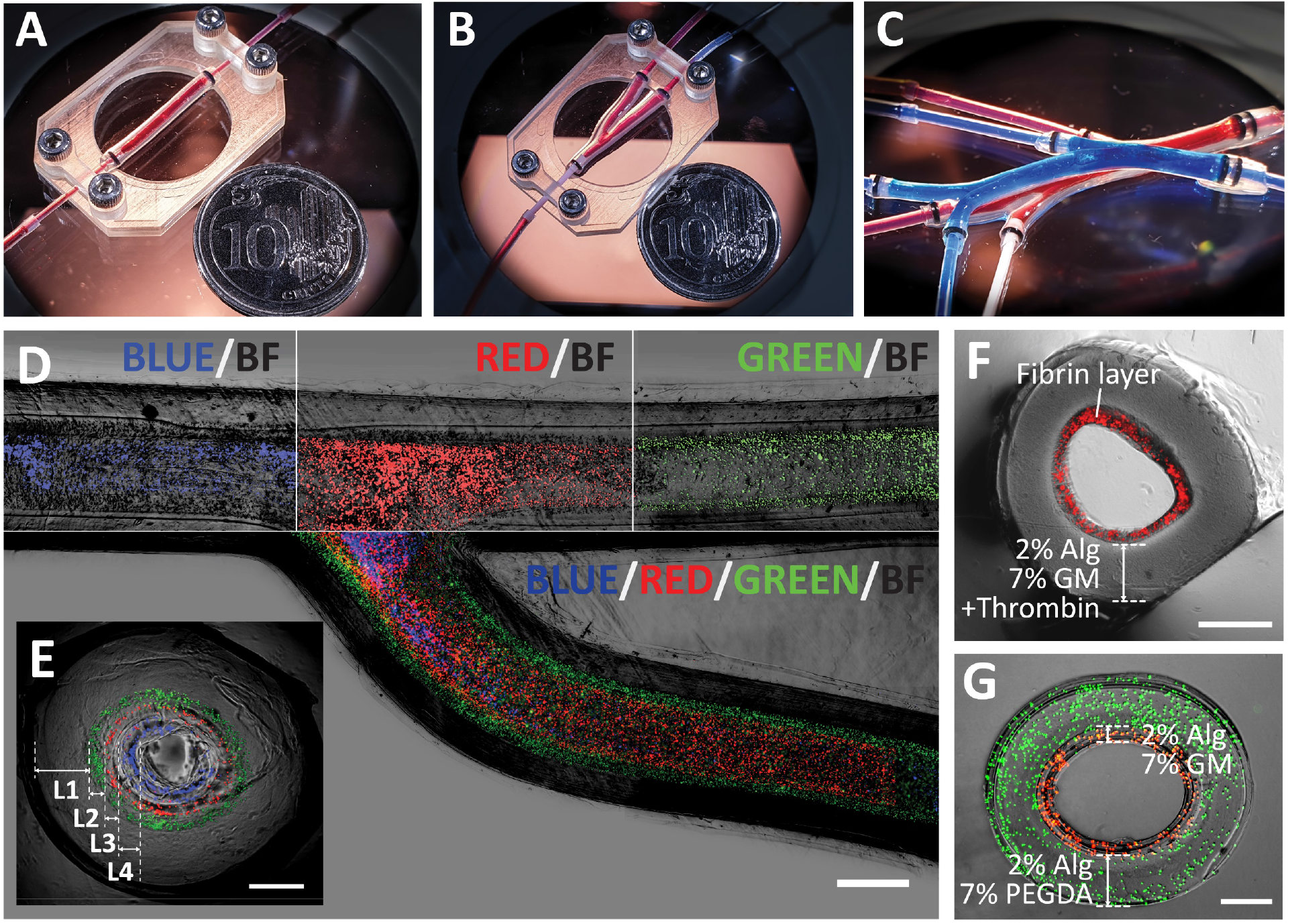
Freestanding vascular constructs recapitulating anatomically-relevant geometries and multilayered structure of a blood vessel. Photographs of **A)** straight vascular construct, **B)** bifurcating vascular construct, and **C)** vascular constructs with multiple branches (where two vasculature constructs are overlaid). The vascular constructs were perfused with red or blue dyes for visualization. Micrograph of a bifurcating vascular construct consisting of four layers (as depicted by the encapsulated fluorescent beads of different colors) shown in **D)** top view and **E)** cross-sectional view. Cross-sectional micrograph of multilayered vascular construct consisting of **F)** Alg-GM in the outer layer and fibrin hydrogel in the inner layer, and **G)** Alg-PEGDA in the outer layer and Alg-GM in the inner layer. Scale bars: **D)** 1 mm, **E)** 500 µm, **F-G)** 500 µm.

Another benefit of multilayered architecture is the ability to tune the mechanical properties of specific layers. In general, cell-responsive hydrogels such as GelMA, fibrin, and ColMA are soft (elastic modulus < 100 kPa), making them challenging to handle and susceptible to unintentional breakage. Furthermore, crosslinked alginate degrades and loses its mechanical integrity after an extended period in cell culture media [56]. The multilayer design allows the blending of stiff and stable hydrogels (*i.e*., PEGDA) in the outer layer to increase mechanical stability while using a blend of hydrogels that support the growth of relevant cells for the inner layer (**Fig. 4G**; the inner layer of Alg-GM and the outer layer of Alg-PEGDA). The ability to tune the mechanical properties of selected layers in the vascular constructs increases the possibility of mimicking the wide-ranging blood vessels (*e.g*., arteries, veins, diseased arteries) found *in vivo* with differing mechanical properties (elastic modulus: 0.1 – 1.5 MPa). These demonstration highlights the versatility of our fabrication technique to create anatomically-relevant vascular constructs with tunable mechanical and biochemical properties achieved through the sequential assembly of multiple layers of bioinks.

### 3.5. Incorporation of vascular cells in a biomimetic configuration

With the success of fabricating multilayer vasculatures with different materials, we then demonstrated the ability to incorporate relevant cell types in our fabricated hydrogel vascular constructs. In particular, SMCs and HUVECs were cocultured as concentric layers to mimic the architectures of the media and intima layers of a blood vessel. To enhance cell spreading within and cell attachment on the hydrogel, we enriched the bioink (2 % (w/v) alginate, 6.7 % (w/v) GelMA) with 50 µL/mg of fibronectin [57-59]. The configuration of a blood vessel consists of endothelial cells in the innermost layer (*i.e*., intima layer), followed by a layer that is predominantly composed of SMCs embedded in a stromal matrix (*i.e*., tunica media). To mimic this tissue configuration, we first suspended the SMCs in the bioink precursor and fabricated the cell-laden vascular construct (**Fig. 5A-B**). After fabrication, the vascular construct was maintained under constant perfusion with a peristaltic pump for seven days (**Fig. S3**). After seven days, HUVECs were seeded on the lumen surface of the hydrogel vascular construct and were maintained under perfusion for three days (**Fig. 5C-D**). After ten (cumulative) days of cell culture, we observed a multilayer configuration analogous to a blood vessel. The SMCs (as shown by phenotypic marker α-SMA) proliferated and migrated within the hydrogel matrix (**Fig. 5E**). The SMCs were mostly aggregated near the surface of the lumen, close to the source of medium flow. The HUVECs were able to form a confluent monolayer in the lumen surface, on top of the SMCs, as indicated by CD31 expression (**Fig. 5E**). The HUVECs in our fabricated vascular construct showed cell-cell adhesion as indicated by vascular endothelial-cadherin (VE-cadherin) expression (**Fig. 5F**). Next, to highlight the uniqueness of our fabrication technique, we fabricated a cell-laden, bifurcating vascular construct incorporated with SMCs and HUVECs (**Fig. 5G**). To the best of our knowledge, such bifurcating, freestanding, multilayered and perfusable vascular constructs incorporating cells has never been demonstrated. Lastly, we assessed the inflammatory response of the HUVECs in our fabricated vascular construct by introducing inflammatory cytokine, tumor necrosis factor-alpha (TNF-α), followed by monocyte adhesion assay. We observed differential monocyte (U937) adhesion in the activated vessel wall, indicating functionality of the HUVECs in the fabricated vascular construct (**Fig. 5H, 5J**). Overall, we demonstrated the incorporation of relevant cell types arranged in multilayered and bifurcating configurations similar to native blood vessels. Importantly, we demonstrated phenotypical expression of relevant cell types and a functional endothelium capable of mimicking inflammatory response. These demonstrations highlight the unique advantage of our fabrication technique and its potential in modeling biomimetic disease models (*i.e*., atherosclerosis, coronary artery bifurcation disease (CABD), bifurcating lesions) [60, 61] that can further our understanding of CVDs, leading to the development of better therapeutic interventions.

**Figure 5.**
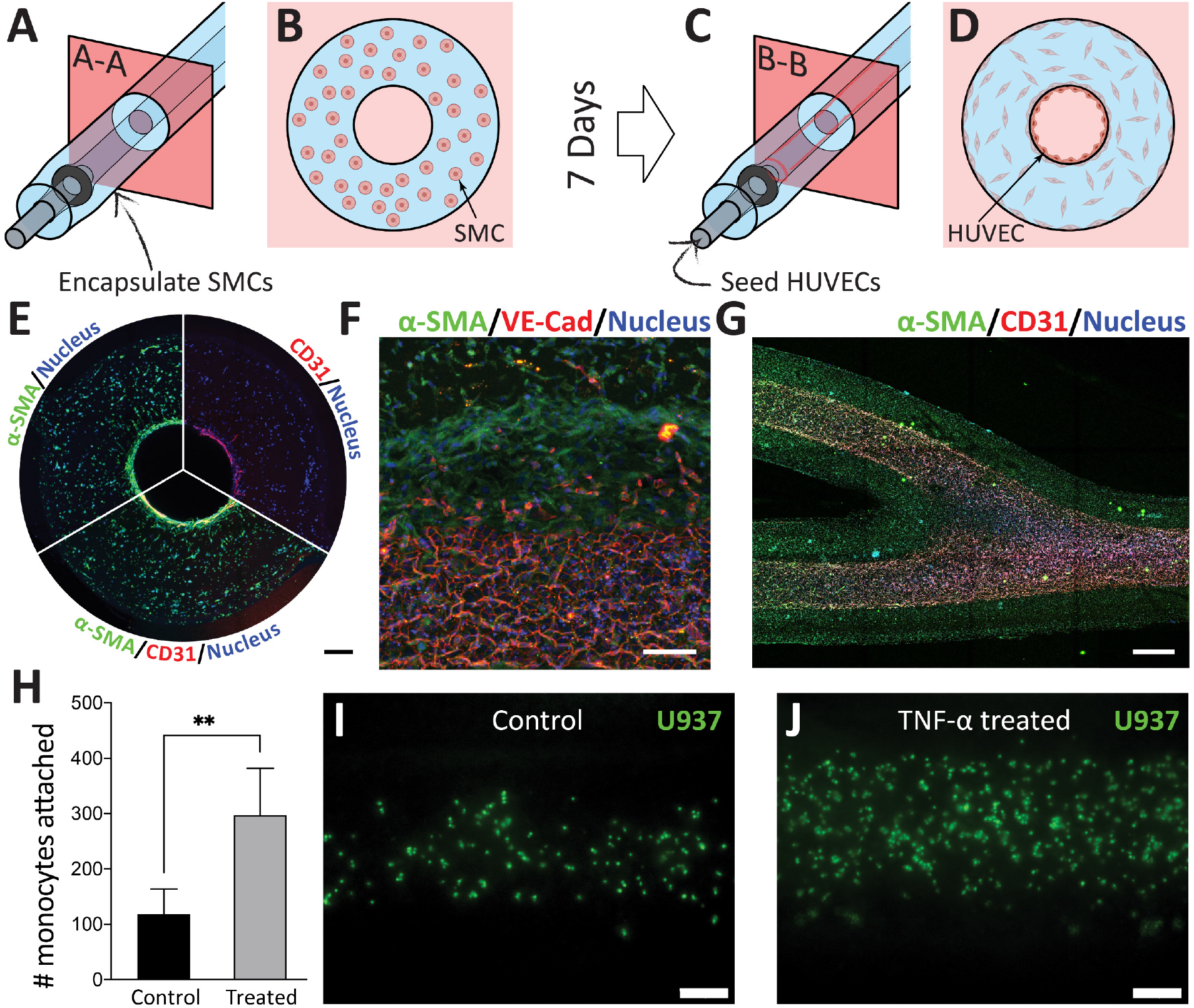
Vascular constructs with encapsulated and surface-cultured cells. **A, B)** Illustration of the encapsulation of SMC in the vascular construct. **C, D**) Illustration of the 2D seeding of HUVECs in the vessel lumen on Day 7. **E)** Cross-sectional confocal micrograph of the vascular construct highlighting the multilayer configuration after perfusion culture on Day 10. **F)** Confocal micrograph highlighting the tight junction present in the confluent layer of HUVECs. **G)** Confocal micrograph of bifurcating vascular construct. **H)** Bar graph showing the adhesion of the monocyte to the fabricated vasculature. **I, J)** Fluorescence micrographs showing the adhered U937 on the lumen surface. Scale bars: **E)** 200 µm, **F)** 100 µm, **G)** 500 µm, **I, J)** 100 µm.

### 3.6. Models for cyclical mechanical loading and deployment of percutaneous coronary interventions

We have shown the main benefit of our proposed fabrication technique, namely (1) the ability to fabricate vascular construct with anatomical accuracy (*i.e*., freestanding, multilayered, and branching) incorporated with relevant vascular cells and (2) the ability to accommodate a wide array of bioinks to achieve cell compatibility and desired mechanical properties. Lastly, we demonstrated the additional biomimicry capabilities of our fabrication technique: cyclical mechanical loading and deployment of percutaneous coronary interventions (PCIs). For instance, coronary arteries are not stationary but experience cyclical stretching as the heart expands and contracts [62]. The cyclical stretching and relaxing of the vascular construct inevitably affect hemodynamics and have been shown to play key roles in developing cardiovascular diseases [63-66]. Organ-on-a-chip systems have attempted to reconstitute physiologically relevant motions [30, 67, 68], but the achievable motions are limited to those predesigned with the device. As our fabricated vascular constructs were freestanding and compliant, they can be fitted to external surfaces that offer the motions of interest. As a demonstration, we placed the fabricated vasculature to an expandable balloon and subjected it to cyclical inflation and deflation (**Fig. 6A-B, Movie S7**). We then performed particle image velocimetry (PIV) to visualize the changes in flow profile when the vascular construct was stretched and relaxed. As expected, the stretching and relaxing of the vascular construct drastically affected the fluid flow profile through a bifurcating vascular construct (**Movie S8**). The flow profile was laminar in the relaxed state with a maximum velocity of 1.6 mm/s (**Fig. 6C**). However, the flow profile was partially chaotic at the stretched state, and the maximum velocity increased to 3.5 mm/s, which was more than doubled in comparison to the relaxed state (**Fig. 6D**). An *in vitro* model that can mimic cyclical stretching of coronary arteries is advantageous to further our understanding of disease progression in the context of hemodynamics. Additionally, our proposed technique enabled the fabrication of vascular constructs with constriction, mimicking blood vessels in the diseased state (*i.e*., intimal hyperplasia, atherosclerosis, cardiovascular stenosis) (**Fig. 6E**). The fabrication of constricted vascular constructs was accomplished by modifying the CAD file for the PEGDA porous molds. Importantly, these freestanding vascular constructs with constrictions were amenable to PCI deployment. We subjected the vascular constructs to drug-eluting balloons (**Fig. 6F**) and drug-eluting stents (**Fig. 6G, Movie S8**). Crucially, existing approaches (*e.g*., co-axial printing, sacrificial molding) do not permit designing and fabricating local anomalies within the freestanding vasculatures, and our methods offer an unprecedented route to realize the unique design of vasculatures. With the incorporation of relevant cells, we envisage that such uniquely designed vasculatures can serve as preclinical models to evaluate and test future PCIs.

**Figure 6.**
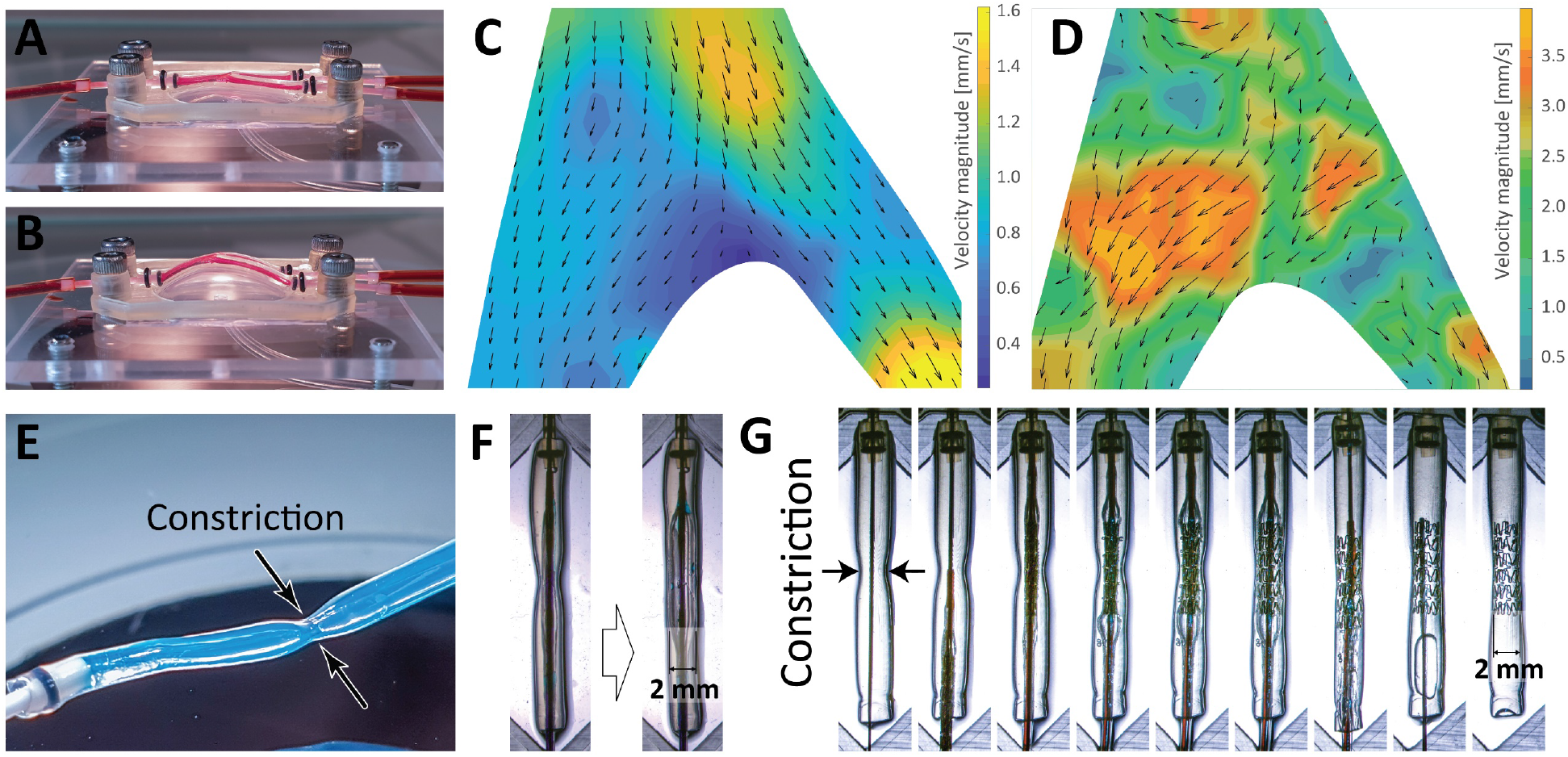
Vascular constructs toward physiologically relevant testbed for vascular disease. Photographs of bifurcating vascular construct under cyclical mechanical loading: **A)** relaxed and **B)** stretched states. PIV analysis of the flow within the bifurcating vascular construct at **C)** relaxed state, and **D)** stretched state. Deployment of percutaneous coronary interventions (PCI): **E)** Photograph of the vascular construct with a constriction. **F)** Micrographs of vascular construct being inflated with a 2-mm angioplasty balloon. **G)** Timelapse micrographs showing the deployment of drug-eluting stent in the vascular construct with a constriction.

## 4. Conclusion

In this research, we have developed a fabrication technique capable of fabricating cell-laden vascular models with (1) high anatomical relevance (*i.e*., freestanding, branching, multilayered, perfusable) and (2) tailored configurations of cells and biomaterials. Importantly, relevant cells (*i.e*., SMCs and HUVECs) were successfully integrated and maintained for 10-day culture under constant perfusion. The cell-laden vascular constructs demonstrated a pronounced increase in monocyte adhesion when tumor necrosis factor-alpha (TNF-α) was introduced, highlighting a potential disease model for mechanistic studies of CVDs (*i.e*., atherosclerosis). The fabricated vasculatures may recapitulate physiologically relevant conditions relevant to CVDs, including (1) vascular geometry (*i.e*., constrictions, bifurcation) and (2) mechanical motion (*i.e*. cyclical motion), which is also amenable to simulate insertion of real medical devices (*e.g*., drug-eluting balloons and drug-eluting stents). Such demonstrations pave routes of employing the fabricated vascular constructs for the model of vascular disease as well as the evaluation of therapeutic intervention. We believe that the versatility and multifaceted possibilities of our fabrication technique will eventually lead to biofabrication of patient-specific vasculatures to benefit future research in mechanistic understanding of CVD progression and the development of therapeutics.

## Acknowledgment

T.C. thanks Ministry of Education (MOE), Singapore, for awarding the President’s Graduate Fellowship. J.J.N. acknowledges Abbott for providing the drug-eluting stents and balloons. M.H. acknowledges Agency for Science, Technology and Research (A*STAR) (A*STAR-AMED joint grant, A19B9b0067) and MOE, Singapore (Academic Research Fund (AcRF) Tier 2, MOE2019-T2-2-192) for the project funding. Y.C.T. acknowledges MOE, Singapore (Tier 1, R-397-000-298-114), Australian Research Council (ARC) Future Fellowship (FT180100157) and Discovery Project (DP200101658).

