## Supplementary Information for "Biomimetic Vasculatures by 3D-Printed Porous Molds"

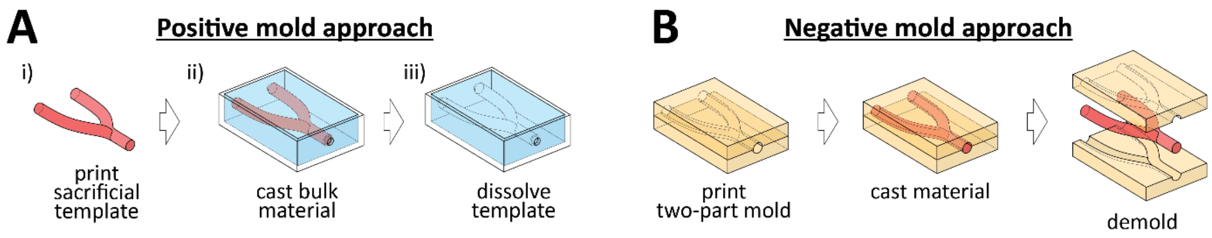


**Figure S1.** Illustrations of **A)** positive mold approach and **B)** negative mold approach for the fabrication of vasculature-inspired structures.

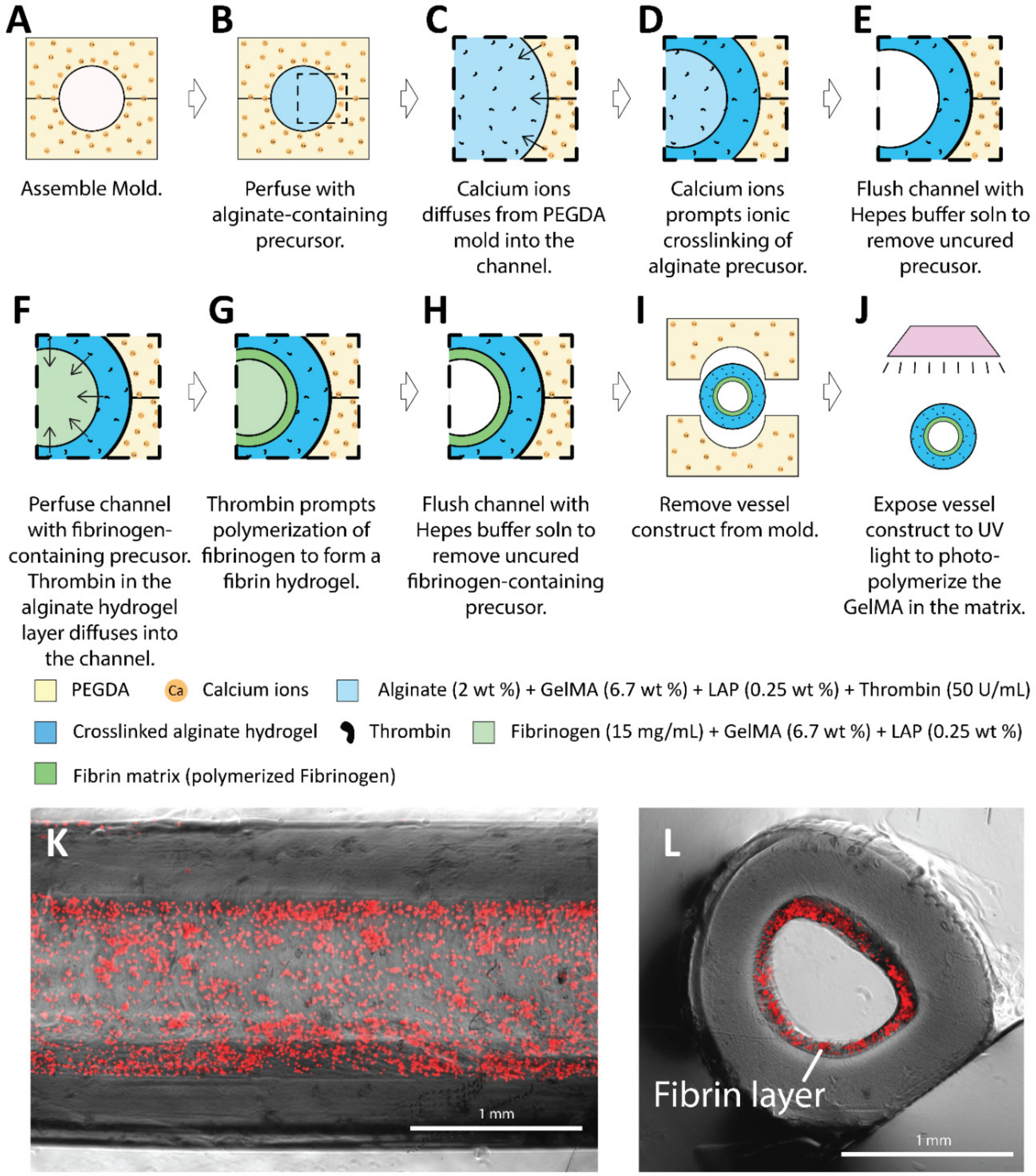


**Figure S2.** Fabrication of the two-layer vascular constructs with a Alg-GM outer layer and a fibrin inner layer. **A-E)** Step-by-step description of the fabrication of outer Alg-GM layer. **F-J)** Step-by-step description of the fabrication of the inner fibrin layer followed by photopolymerization. Micrographs showing the **K)** top view and **L)** cross-sectional view of the vascular construct with fibrin inner layer (encapsulated with red fluorescence beads).

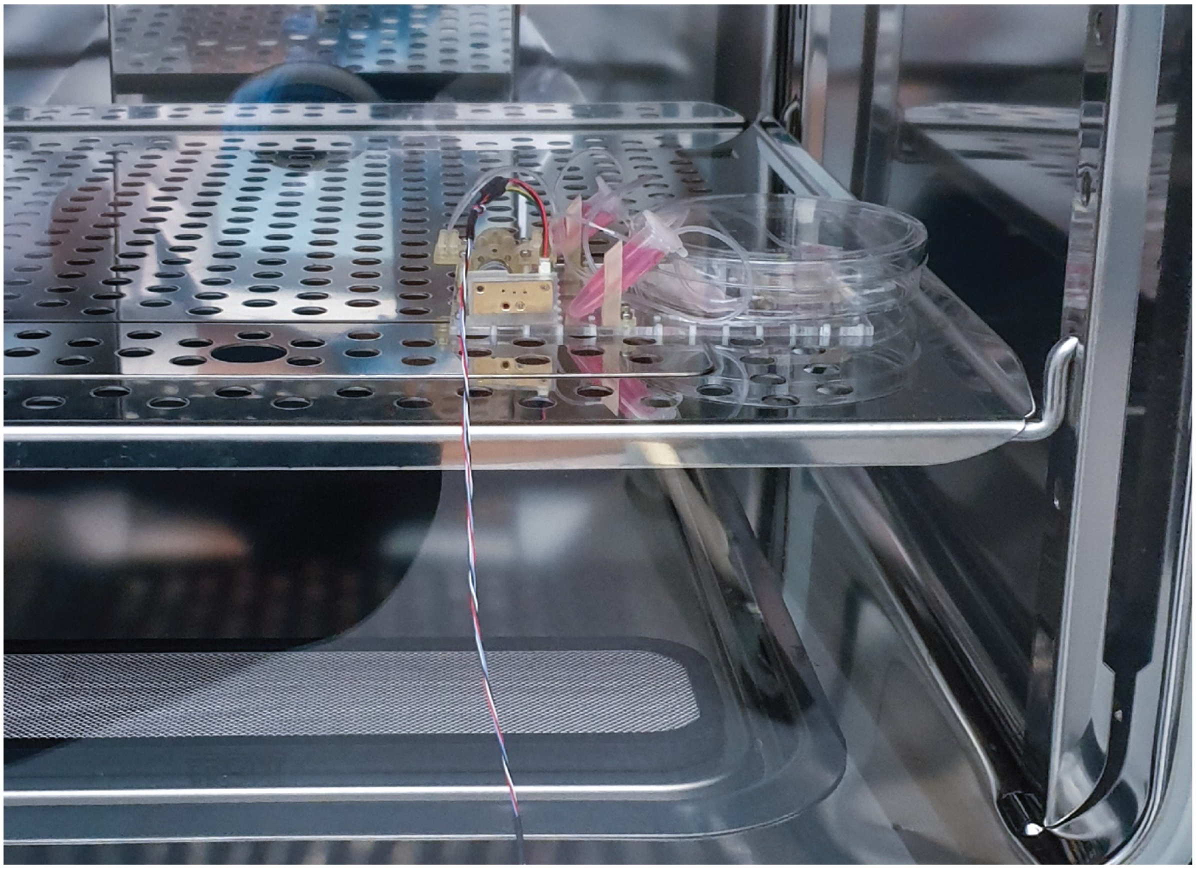

**Figure S3.** Perfusion culture of the vascular construct using a DIY peristaltic pump.

**Table S1.** Summary of existing available methods for 3D printing of biomimetic vascular constructs and their respective capabilities

| *Fabrication method* | *Freestanding* | *Complex architecture* | *Multilayer organization* | *Perfusion culture* | *Cyclic stretching* | *Percutaneous coronary interventions* * | *Ref.* |
|---|---|---|---|---|---|---|---|
| *Sacrificial molding* | No | Multi-scale, hierarchical networks | n.d. | Yes | n.d. | n.d. | [24-30] |
| *Direct ink writing (DIW); Scaffold-free bioprinting* | Yes | Branching networks | 2 layers | Yes | n.d. | n.d. | [16-18] |
| *Embedded 3D printing* | Yes | Multi-scale, hierarchical networks | 2 layers | Yes | n.d. | n.d. | [19-22] |
| *Light-assisted 3D printing (e.g., SLA, DLP)* | Yes | Multi-scale, hierarchical networks | n.d. | Yes | n.d. | n.d. | [23] |
| *Coaxial bioprinting* | Yes | Not suitable | 2 layers | Yes | n.d. | n.d. | [31-34] |
| *Molding by 3D-printed porous mold* | Yes | Multi-scale, hierarchical networks; constriction | 4 layers | Yes | Yes | Yes | This work |

**n.d.**, no data/not demonstrated; * *Inserson of drug-eluting balloons and stents*

**Table S2.** Advantages and limitations of available methods for 3D printing of biomimetic vascular constructs.

| *Fabrication method* | *Materials* | *Size scale* | *Advantages* | *Limitations* | *Ref.* |
|---|---|---|---|---|---|
| *Sacrificial Molding* | **Sacrificial Material:** carbohydrate powders; F127 pluronic acid; agarose; PVA **Bulk Material:** PDMS; PCL; PEGDA; Agarose; Silk Fiboin; Fibrin; GelMA; Gelatin | **LD:** 100 – 1000 µm | Wide selection of bioactive agents for bulk material<br><br>Facilitates the fabrication of complex, multiscale networks | Sacrificial mold must be removable<br><br>Additional steps required to evacuate sacrificial materials | [24-30] |
| *Direct ink writing (DIW); Scaffold-free bioprinting* | Nanoengineered ionic covalent entanglement (NICE); PEGDA + alginate; agarose | **LD:** ~0.9 – 22 mm **WT:** >300 µm | Easy setup<br><br>Cost-effective<br><br>Suitable for large scale (>1 mm) models | Require the control of printed bioinks (*e.g.*, yield-stress fluids, thickening agents, photopolymerization).<br><br>Large attainable lumen diameter (> 900 µm) | [16-18] |
| *Embedded 3D printing* | **Suspension media:** Granular gel; polyacrylic acid; hyaluronic acid **Bioink:** Bacterial cellulose; PVA; fibrinogen; collagen; Matrigel; decellularized ECM | **WT:** >100 µm **LD:** >200 µm | Bioink need not to be yield stress fluid.<br><br>Wide selction of printable bioinks | Require suspension media made up of yield stress fluid<br><br>Requires removal of suspension media to achieve freestanding constructs<br><br>Entire printing process can be slow | [19-22] |
| *Light-assisted 3D printing (e.g., SLA, DLP)* | PEGDA; PEGDA + GelMA; | **LD:** ~ 300 µm | High fabrication speed (compared to embedded 3DP and DIW)<br><br>Good spatial resolution | Limited photocurable bioinks available | [23] |
| *Coaxial bioprinting* | Alginate-containing hydrogel (*e.g.,* Alginate + decellularized vascular ECM, Alginate + GelMA, Alginate + PEGTA) | **LD:** ~500 – 2000 µm **WT:** ~200 – 1000 µm | Facilitates the fabrication of multilayered vascular constructs with relevant vascular cells<br><br>Fast print speed | Require bioink that contains alginate<br><br>Branching networks not achievable<br><br>Requires the assembly of concentric, multilayered nozzle | [31-34] |
| *Molding by 3D-printed porous mold* | Alginate-containing hydrogel (*e.g.,* Alginate + GelMA, Alginate + PEGDA); Fibrin | **WT:** >200 µm **LD:** >500 µm | Multi-branching, multilaylared vascular constructs (4 layers) with relevant cells<br><br>Circumvents the challenges in direct printing and handling of bioinks | Require bioink blend that contains alginate | This work |

**LD**, lumen diameter; **WT**, wall thickness, **n.d.**, no data/not demonstrated; PEGDA, poly(ethylene glycol) diacrylate; GelMA, gelatin methacryloyl; PVA, poly(vinyl alcohol); PEGTA, poly(-ethylene glycol)-tetra-acrylate; PCL, polycaprolactone

**Table S3.** Summary of staining protocols and reagents

| **Antibody or stain** | **Source** | **Catalog#** | **Host Species & Reactivity** | **Concentration** |
|---|---|---|---|---|
| **CD31** | Cell Signaling Technologies | 3528 | Mouse anti-human | 1:200 |
| **VE-Cadherin** | Cell Signaling Technologies | 2500 | Rabbit anti-human | 1:200 |
| **Anti-alpha smooth muscle Actin antibody** | Abcam | ab5694 | Rabbit anti-human | 1:200 |
| **Anti-alpha smooth muscle Actin antibody** | Abcam | ab7817 | Mouse anti human | 1:200 |
| **Sytox-green** | Invitrogen | S7020 | N/A | 1:10000 |
| **CellTracker-Green** | Invitrogen | C2102 | N/A | 1:1000 |
| **Alexa Fluor 647** | Life Technologies | A-31573 | Donkey anti-Rabbit | 1:500 |
| **Alexa Fluor 647** | Life Technologies | A-31571 | Donkey anti-mouse | 1:500 |
| **Alexa Fluor 555** | Life Technologies | A-31572 | Donkey anti-Rabbit | 1:500 |
| **Alexa Fluor 555** | Life Technologies | A-31570 | Donkey anti-mouse | 1:500 |
